# The Mimivirus L375 Nudix enzyme hydrolyzes the 5’ mRNA cap

**DOI:** 10.1101/2021.01.11.426193

**Authors:** Grace Kago, Susan Parrish

## Abstract

The giant Mimivirus is a member of the nucleocytoplasmic large DNA viruses (NCLDV), a group of diverse viruses that contain double-stranded DNA (dsDNA) genomes that replicate primarily in eukaryotic hosts. Two members of the NCLDV, Vaccinia Virus (VACV) and African Swine Fever Virus (ASFV), both synthesize Nudix enzymes that have been shown to decap mRNA, a process thought to accelerate viral and host mRNA turnover and promote the shutoff of host protein synthesis. Mimivirus encodes two Nudix enzymes in its genome, denoted as L375 and L534. Importantly, L375 exhibits sequence similarity to ASFV-DP and eukaryotic Dcp2, two Nudix enzymes shown to possess mRNA decapping activity. In this work, we demonstrate that recombinant Mimivirus L375 cleaves the 5’ m^7^GpppN mRNA cap, releasing m^7^GDP as a product. L375 did not significantly cleave mRNAs containing an unmethylated 5’GpppN cap, indicating that this enzyme specifically hydrolyzes methylated-capped transcripts. A point mutation in the L375 Nudix motif completely eliminated cap hydrolysis, showing that decapping activity is dependent on this motif. Addition of methylated cap derivatives or uncapped RNA inhibited L375 decapping activity, suggesting that L375 recognizes its substrate through interaction with both the mRNA cap and RNA body.

## Introduction

The giant Mimivirus, which infects *Acanthamoeba* species, possesses a 1.2 Mb genome encoding over 900 proteins [1–3]. Interestingly, Mimivirus rivals some small bacterial species with respect to physical and genome size, and was the first virus identified to encode some of its own translational components [1–3]. Mimivirus is a member of the nucleocytoplasmic large DNA viruses (NCLDV), a group that currently includes seven viral families: *Poxviridae, Asfarviridae, Iridoviridae, Phycodnaviridae, Mimiviridae*, *Ascoviridae,* and *Marseilleviridae* [4–8]. While many members of the NCLDV share a subset of conserved genes, the evolutionary origins and relationships of the NCLDV remain controversial [4–7, 9].

The Nudix hydrolase motif is a conserved amino acid sequence found in a diverse group of enzymes that typically cleave *nu*cleoside *di*phosphates linked to another moiety *X* [10]. Nudix enzymes are nearly universal, found in prokaryotes, eukaryotes, and some viruses [4, 5, 11, 12]. Two NCLDV families, *Poxviridae* and *Asfarviridae*, encode Nudix enymes that possess intrinsic mRNA decapping activity, a process that leads to mRNA degradation and subsequent inhibition of gene expression [13–15].

The prototypic poxvirus, Vaccinia Virus (VACV), encodes two Nudix enzymes in its genome termed D9 and D10. The D9R (VACV-WR_114) and D10R (VACV-WR_115) genes lie adjacent to each other in the genome and encode proteins that share ~20% amino acid identity but are expressed at different times during infection [16–18]. Prior genetic studies showed that over-expression of either D9R or D10R resulted in accelerated turnover of mRNAs containing 5’ m^7^GpppN caps, a structural feature of both VACV and eukaryotic host mRNAs [17]. While deletion of D9R did not cause any obvious defects, deletion or inactivation of D10R resulted in persistence of viral and host mRNAs and a delay in the shutoff of host protein synthesis [17–19]. Subsequent biochemical experiments confirmed that both D9 and D10 cleave the mRNA cap, releasing m^7^GDP as a product [13, 14]. Together, these data suggest that VACV D9 and D10 decap viral and host mRNAs to facilitate mRNA turnover and the shutoff of host protein synthesis, thereby promoting viral infection.

The sole member of *Asfarviridae*, African Swine Fever Virus (ASFV), contains a gene (termed g5R in strain Malawi and D250 in strain Ba71V) that encodes the Nudix enzyme denoted as ASFV-DP [20, 21]. Although ASFV-DP does not share significant sequence similarity to VACV D9 or D10, it does exhibit sequence similarity to Dcp2, an mRNA decapping enzyme found in yeasts and mammals, along with other multicellular eukaryotes [12, 21–26]. ASFV-DP was shown to cleave a broad range of substrates, including diphosphoinositol polyphosphates, GTP, and the 5’ m^7^GpppN cap when attached to an RNA moiety [15, 20]. Subsequent *in vivo* studies revealed that over-expression of ASFV-DP increases viral and host mRNA turnover, supporting the idea that ASFV-DP mediates mRNA decapping and destabilization during infection [21]

The Mimivirus genome encodes two putative Nudix enzymes in its genome, termed L375 (NCBI ID: YP_003986880) and L534 (NCBI ID: YP_003987047). Of the characterized viral Nudix enzymes, L375 is most similar to the ASFV-DP mRNA decapping enzyme, sharing ~21% amino acid identity. In contrast, L534 does not exhibit significant similarity to any of the known viral or eukaryotic mRNA decapping enzymes, instead showing sequence similarity to uncharacterized Nudix enzymes of other giant viruses, bacteria, and single-celled eukaryotes. Given the sequence similarity between L375 and ASFV-DP, we chose to focus this work exclusively on Mimivirus L375.

To determine if Mimivirus L375 can cleave the mRNA cap, a recombinant version of this protein was expressed in bacteria and purified by affinity chromatography. Recombinant L375 cleaved the 5’ m^7^GpppN mRNA cap, releasing m^7^GDP as a product, a reaction dependent an intact Nudix motif. Importantly, L375 did not significantly cleave unmethylated GpppG capped mRNAs, suggesting that L375 specifically recognizes the methylated mRNA cap structure. Addition of uncapped RNA to the reaction significantly inhibited L375 cap cleavage activity. Furthermore, L375 decapping activity was reduced in the presence of certain methylated cap derivatives, suggesting that L375 uses both the RNA and cap moieties to locate target substrates.

## Materials and methods

### Plasmid design

Mimivirus L375 appended with a C-terminal 10X histidine tag was generated by GeneArt (Life Technologies) for codon-optimized expression in *Escherichia coli.* The synthetic L375 gene was then amplified by the polymerase chain reaction (PCR) using the oligonucleotide primers: 5’-ATG GAA TAT GAA ACC AAC TTT CGC AAA AAA CAC ATT TG and 5’-GCG CGC AAG CTT TTA GTG ATG ATG GTG GTG ATG GTG ATG ATG ATG. The resulting PCR product was ligated into the pMal-c2x protein expression plasmid (New England Biolabs) adjacent to the *malE* gene to generate the pMAL-c2x-*malE*-L375-his_10_ plasmid encoding maltose binding protein (MBP) fusion protein MBP-L375-HIS_10_. A targeted mutation in the Nudix motif was introduced through use of the QuikChange site-directed mutagenesis kit (Agilent Technologies) to produce a plasmid encoding L375 (E258Q).

### Synthesis and purification of recombinant L375 protein

Wild-type or mutated pMAL-c2x-*malE*-L375-his_10_ plasmids were transformed into *Escherichia coli* strain BL21 (EMD Millipore) for subsequent growth in LB broth supplemented with 50 μg/ml carbenicillin and 0.2% (w/v) glucose. MBP-L375-HIS_10_ expression was induced with 0.15 mM isopropyl β-D-1 thiogalactopyranoside (IPTG) followed by growth at 28 °C. After 4 h of induction, cell lysates were produced using sonication. The recombinant protein was sequentially purified using an amylose column (New England Biolabs) followed by a nickel-nitrilotriacetic acid column (Qiagen) and then dialyzed against buffer comprised of 10 mM Tris-HCl pH 7.5, 100 mM NaCl, 10% glycerol, 1 mM DTT, and 2 mM Mg acetate [27]. Recombinant VACV MBP-D10 was produced and purified by affinity column chromatography as detailed in Parrish et al. [13].

### Synthesis of RNA substrate

The MEGAshortscript kit (Life Technologies) and pTRI-β-actin-human template (Life Technologies) were used to *in vitro* transcribe a 309-nt actin RNA transcript that was subsequently cap-labeled using recombinant VACV guanylyltransferase/guanine-7-methyltransferase (Epicentre Biotechnologies) in conjunction with capping buffer (50 mM Tris-HCl pH 8.0, 6 mM KCl, 1.25 mM DTT, 1.25 MgCl_2_), 0.132 μM [α^32^P] GTP, and 0.1 mM *S*-adenosylmethionine [28]. The cap-labeled RNA was then purified from unincorporated nucleotides by using a ProbeQuant G-50 gel filtration column (GE Healthcare).

### RNA decapping assays

0.02 pmol of cap-labeled RNA was incubated with purified recombinant wild-type or mutant MBP-L375-HIS_10_ in the presence of decapping buffer (100 mM K acetate, 10 mM Tris-HCl pH 7.5, 2 mM MgCl_2_, 0.5 mM MnCl_2_, and 2 mM DTT) in a total volume of 15 μl for 30 min at 37 °C [27]. 2 μl aliquots of each reaction were resolved on a polyethyleneimine-cellulose thin layer chromatography plate (Sigma-Aldrich) developed in 0.75 M LiCl. UV shadowing was used to detect unlabeled nucleotide standards while autoradiography and PhosphorImager analysis (Molecular Dynamics) were used to visualize radioactive signals.

## Results

### Recombinant Mimivirus L375 exhibits mRNA decapping activity

The observation that Mimivirus L375 harbors a Nudix motif, in conjunction with the sequence similarity observed between L375 and ASFV-DP, suggested that L375 could possess intrinsic mRNA decapping activity to modulate mRNA turnover during infection. To evaluate if recombinant Mimivirus L375 could decap mRNA, a maltose binding protein (MBP)-L375 fusion protein terminated with C-terminal 10X histidine epitope tag (MBP-L375-HIS_10_) was synthesized in *Escherichia coli* and purified through successive amylose and nickel-nitrilotriacetic acid columns. Following separation of the purified recombinant protein through sodium dodecyl sulfate-polyacrylamide gel electrophoresis (SDS/PAGE), the expected ~87–kDa band for MBP-L375-HIS_10_ was observed (Fig 1A).

**Fig 1.**
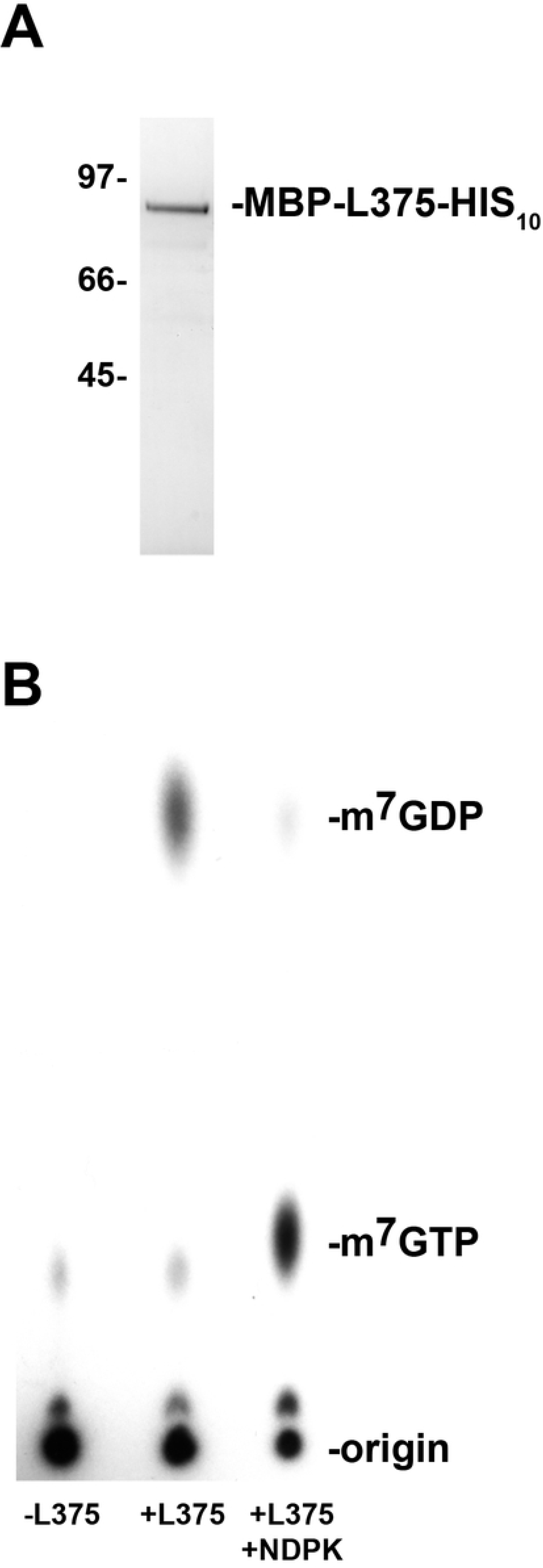
Recombinant Mimivirus L375 hydrolyzes the mRNA cap. (A) An MBP-L375 fusion protein containing a C-terminal 10X histidine epitope tag (MBP-L375-HIS_10_) was synthesized in *Escherichia coli* and purified by affinity chromatography through successive amylose and nickel-nitrilotriacetic acid columns. The purified MBP-L375-HIS_10_ protein was separated by SDS/PAGE and visualized by Coomassie blue staining. The locations of the protein mass standards (in kDa) are labeled on the left side of the gel and the ~87 kDa MBP-L375-HIS_10_ protein is denoted on the right. (B) MBP-L375-HIS_10_ (80 ng) and 0.02 pmol ^32^P-cap-labeled actin RNA were added to decapping buffer and incubated at 37 °C for 30 min. After the incubation, an aliquot of the reaction was treated with 2 U of nucleoside diphosphate kinase (NDPK) in the presence of 1 mM ATP at 37 °C for 30 min to convert nucleoside diphosphates into nucleoside triphosphates. The products of the reaction were separated on PEI-cellulose TLC plates in 0.75 M LiCl and the radioactive signals were visualized by autoradiography. Non-radioactive nucleotide standards were run in parallel and detected by UV shadowing, as indicated on the right.

For the mRNA decapping assays, *in vitro* synthesized 309-nt actin RNA was capped, methylated, and radioactively labeled using recombinant VACV RNA guanylyltransferase/guanine-7-methyltransferase in the presence of [α^32^P] GTP and *S*-adenosylmethionine [28]. The ^32^P-cap-labeled RNA substrate was then combined with MBP-L375-HIS_10_ and the products of the reactions were separated on polyethyleneimine (PEI)-cellulose thin layer chromatography (TLC) plates. Radioactive signals were visualized using either autoradiography or PhophorImager analysis, whereas unlabeled TLC nucleotide standards were detected by UV shadowing.

As expected, in the absence of recombinant L375, the large ^32^P-cap-labeled RNA stayed at the origin of the TLC plate and no major reaction products were detected; the minor faint spot detected may correspond to unincorporated GTP remaining after purification of the cap-labeled RNA (Fig 1B). When recombinant MBP-L375-HIS_10_ was included in the reaction, a product was released that migrated the same distance as the unlabeled m^7^GDP standard, demonstrating that L375 cleaves the methylated mRNA cap (Fig 1B). After cap hydrolysis, a residual amount of uncleaved cap-labeled RNA was observed at the origin. Since L375 specifically cleaves methylated-capped structures (see below) and methylation by *S*-adenosylmethionine of the cap-labeled RNA is generally incomplete, some intact, unmethylated cap-labeled RNA is expected to remain at the origin.

To confirm that the product released was m^7^GDP, the decapping products were incubated with nucleoside diphosphate kinase (NDPK), an enzyme that phosphorylates nucleoside diphosphates to yield nucleoside triphosphates. After the addition of NDPK, the m^7^GDP product generated by MBP-L375-HIS_10_ shifted in a downward direction to co-migrate with the m^7^GTP standard (Fig 1B).

### Recombinant Mimivirus L375 mRNA decapping activity is dependent on the Nudix motif

The highly conserved Nudix box consists of the signature sequence GX_5_EX_5_[UA]XREX_2_EEXGU (where U indicates either isoleucine, leucine, or valine and X denotes any amino acid), of which the EX_2_EE sequence has been shown to be required for divalent cation binding and catalytic activity [10, 29, 30]. To determine whether the Nudix motif was essential for cap cleavage, a point mutation was created in the essential EX_2_EE active site residues of L375. A mutated version of L375 was synthesized in which the glutamic acid at position 258 was changed to glutamine [L375(E258Q)]. The mutated L375(E258Q) protein was expressed in *Escherichia coli* and then purified by affinity chromatography concurrently with the wild-type recombinant proteins. As previously shown, incubation of wild-type recombinant L375 with the capped mRNA substrate resulted in the hydrolysis of the 5’ cap and release of m^7^GDP (Fig 2). However, when an equal amount of the mutant L375(E258Q) protein was added to the capped RNA substrate, m^7^GDP was not liberated, confirming that the Nudix motif was essential for cap cleavage and that the recombinant protein was directly responsible for the decapping activity observed in the assay (Fig 2).

**Fig 2.**
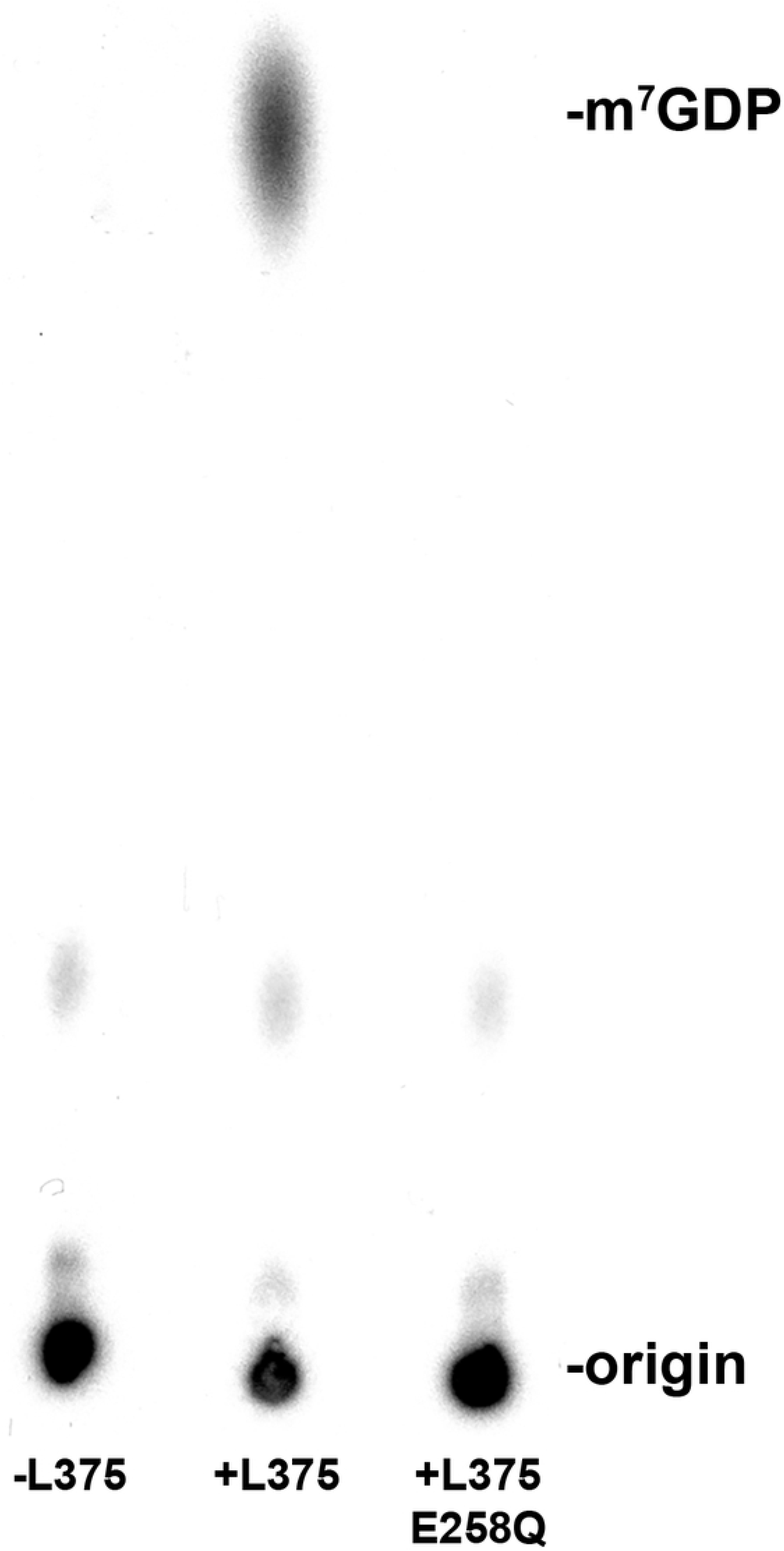
The mRNA decapping activity of recombinant L375 is dependent on an intact Nudix hydrolase motif. The Nudix motif of MBP-L375-HIS_10_ was subjected to site-directed mutagenesis to convert the glutamic acid residue at position 258 into a glutamine residue, thereby producing L375(E258Q). The mutant and wild-type proteins were expressed and purified in parallel as described in Fig 1A and equivalent amounts of the two proteins (50 ng) were added to separate mRNA decapping assays conducted as in Fig 1B.

### Recombinant Mimivirus L375 specifically cleaves methylated cap structures

Since Nudix enzymes can cleave a broad range of substrates, it was important to investigate the specificity of L375 for the 5’ m^7^GpppN mRNA cap [31]. The mRNA cap contains a unique methyl group at position 7 on the guanine base, a structural feature often recognized by proteins that exclusively bind and/or cleave the mRNA cap. To examine the selectivity of L375 for the 5’ m^7^GpppN cap structure, an unmethylated GpppN-capped RNA was synthesized by excluding the *S*-adenosylmethionine methyl donor from the capping reaction, and the unmethylated substrate was subsequently incubated with recombinant L375. Importantly, recombinant L375 was not able to significantly cleave the unmethylated GpppN cap to release GDP, indicating that this enzyme specifically recognizes the methylated mRNA cap (Fig 3). As expected, when equivalent amounts of mutant L375(E258Q) protein was added to the unmethylated cap-labeled substrate, no products were generated.

**Fig 3.**
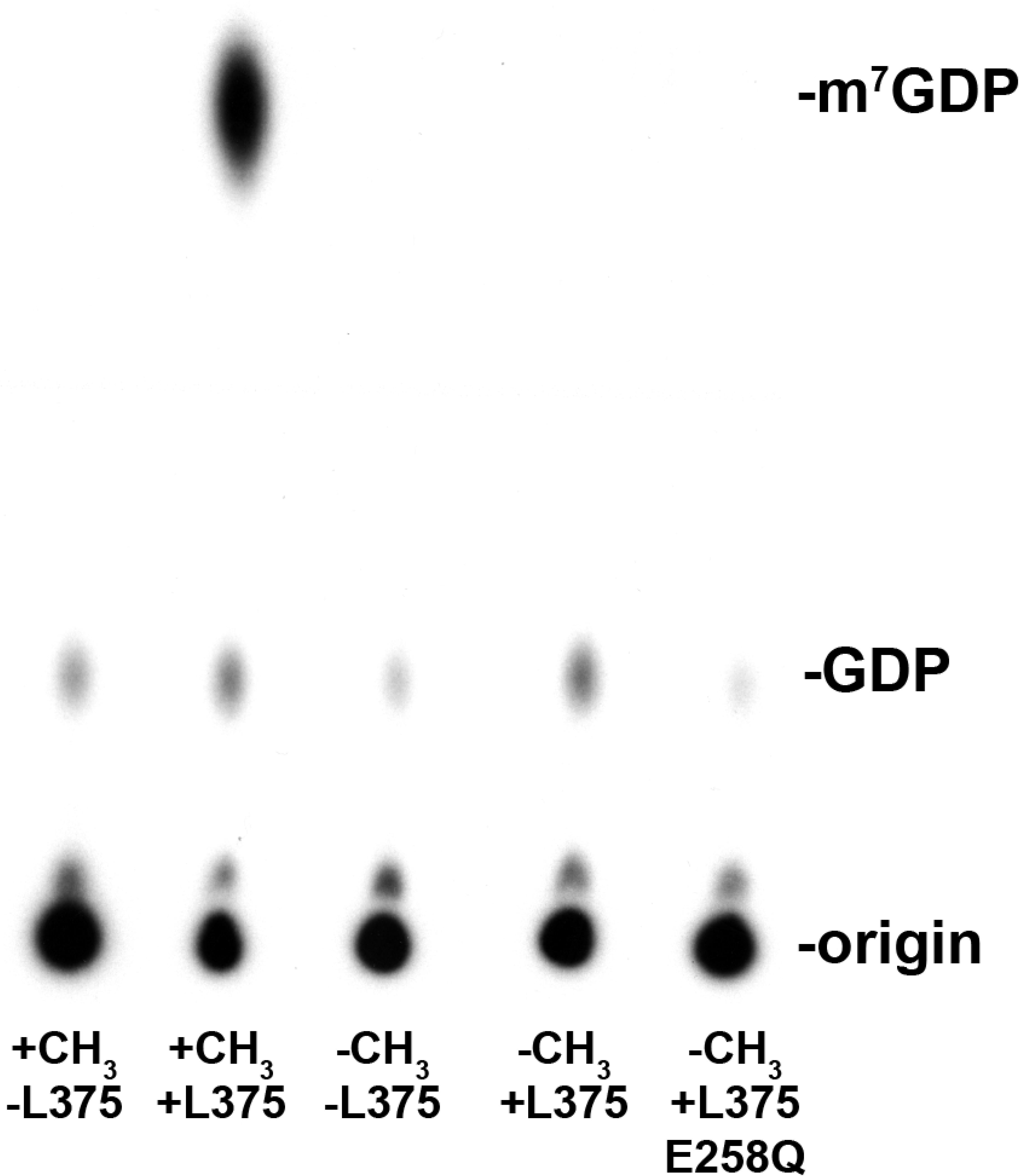
Recombinant L375 specifically cleaves methylated cap structures. (A) The ^32^P-cap-labeled actin 309-nt RNA substrate was synthesized either in the presence (+CH_3_) or absence (−CH_3_) of the methyl donor *S-*adenosylmethionine. 0.02 pmol of methylated or unmethylated RNA substrate was incubated with either 50 ng of recombinant wild-type L375 or mutant L375 (E258Q) in mRNA decapping assays as described in Fig 1B.

### Mimivirus L375 mRNA decapping activity increases with time and enzyme concentration

A time course experiment revealed that the amount of m^7^GDP product released by L375 increased through time, as expected for an enzyme (Fig 4A). Likewise, increasing amounts of L375 enzyme resulted in the release of more m^7^GDP product, until saturation was reached (Fig 4B). As noted for above and similar to other viral mRNA decapping enzymes, a proportion of the substrate remains resistant to cleavage despite high enzyme concentrations (~28%), because some of the ^32^P-cap-labeled RNA remains unmethylated and therefore not a viable substrate. This observation, along with the finding that the presence of uncapped RNA inhibits L375 mRNA decapping activity (see Fig 5 below), hindered our ability to determine the kinetic coefficients of the reaction.

**Fig 4.**
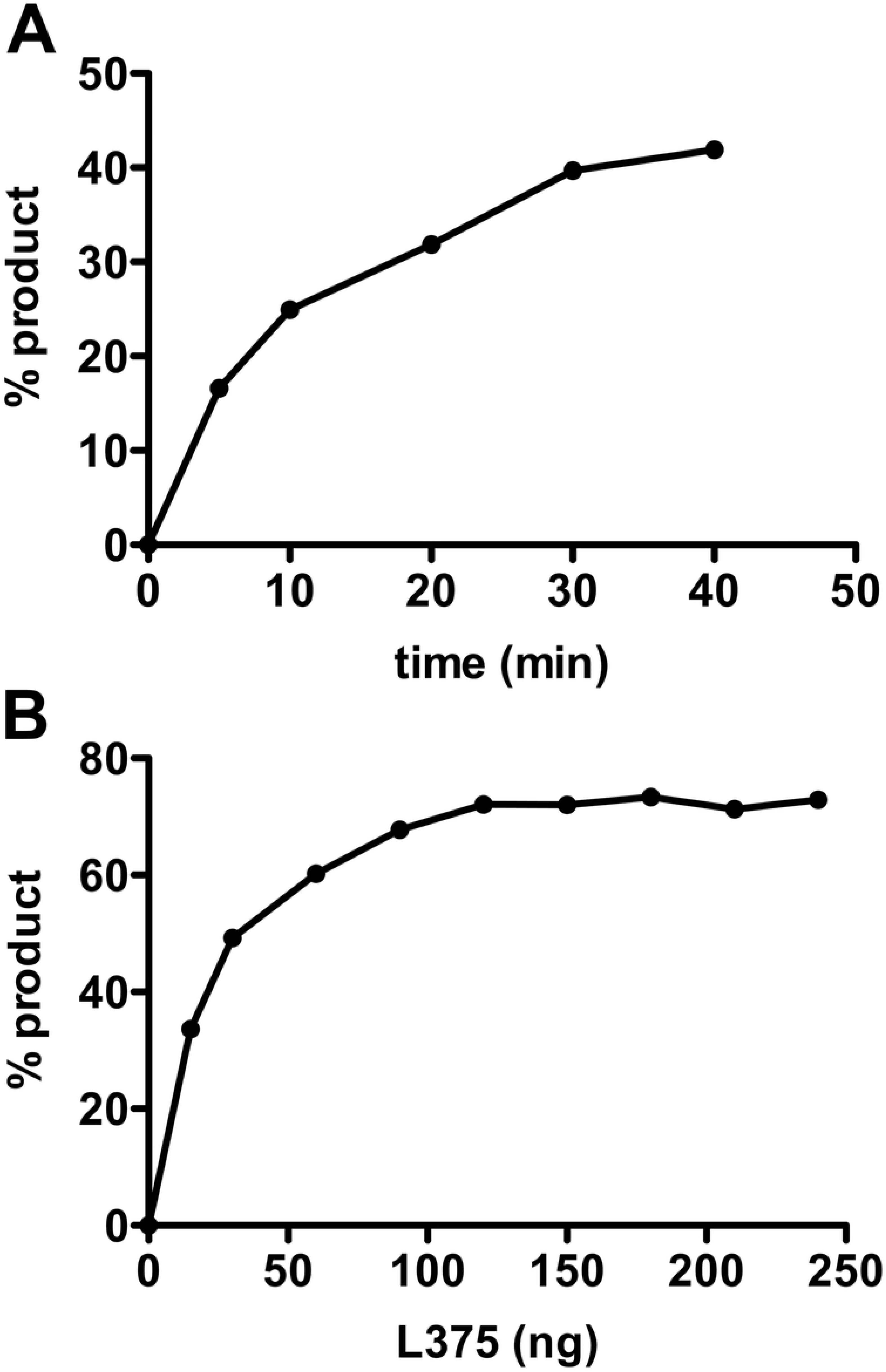
Recombinant L375 mRNA decapping activity increases with time and enzyme concentration. (A) 60 ng of recombinant L375 was incubated with 0.02 pmol ^32^P-cap-labeled actin RNA substrate in decapping buffer at 37 °C for the time indicated on the graph. After separation of the reaction products on PEI-cellulose TLC plates, the percentage of m^7^GDP released was calculated by PhosphorImager analysis. (B) The indicated amount of recombinant L375 was added to 0.02 pmol ^32^P-cap-labeled actin RNA substrate for mRNA decapping assays performed as in Fig 1B. The percentage of m^7^GDP liberated was quantified using a PhosporImager.

**Fig 5.**
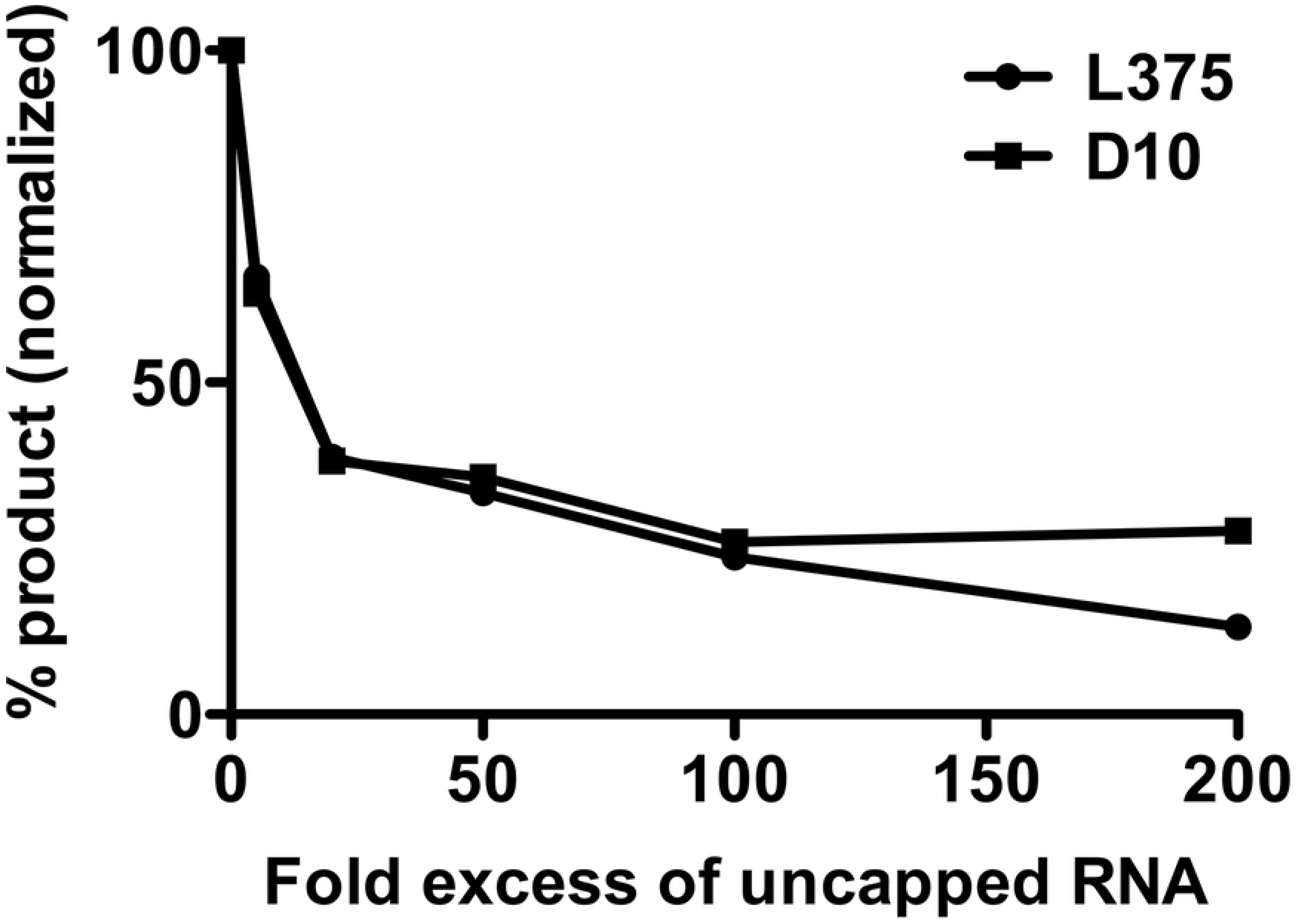
Recombinant L375 mRNA decapping activity is reduced by the addition of uncapped RNA competitor. 80 ng of recombinant L375 or VACV D10 and 0.02 pmol ^32^P-cap-labeled 309-nt actin RNA were incubated with increasing amounts of uncapped, non-radioactive 309-nt actin RNA in mRNA decapping assays conducted as described in Fig 1B. The percentage of m^7^GDP product released was determined by using a PhosphorImager.

### Uncapped RNA inhibits Mimivirus L375 mRNA decapping activity

The viral mRNA decapping enzymes characterized to date have all been shown to be inhibited by the addition of uncapped RNA, suggesting that these enzymes recognize and bind the RNA moiety during substrate identification [13–15]. To evaluate if Mimivirus L375 also interacts with the RNA body during substrate recognition, increasing molar amounts of uncapped RNA were added to the reaction and the percentage of m^7^GDP product released was calculated. Addition of uncapped RNA competitor significantly decreased L375 decapping activity, in a manner almost identical to that observed for VACV D10 (Fig 5). For example, a 20-fold molar excess of competitor RNA reduced both L375 and VACV decapping activity by 62%, suggesting that L375 binds the RNA body during substrate recognition (Fig 5).

### Methylated nucleotides inhibit L375 mRNA decapping activity

It was previously shown that the decapping activity of VACV D9 and D10 was inhibited by m^7^GTP, m^7^GDP, and m^7^GpppG, structures that mimic the mRNA cap and therefore may compete for binding [13, 14]. Conversely, ASFV-DP was not inhibited by any of the three methylated cap analogs, indicating that ASFV-DP may exclusively use the RNA moiety to locate its substrate [15]. To evaluate the effect of methylated nucleotides on L375 substrate cleavage, increasing quantities of m^7^GTP, m^7^GDP, and m^7^GpppG or unmethylated alternatives of these nucleotides were included in the reactions and the amount of product generated was calculated. Interestingly, the most robust inhibition of L375 decapping activity was observed for m^7^GTP, mirroring the results seen for VACV D10 (Figs 6A, 6B, and 6C). m^7^GDP also inhibited L375 cap cleavage, but the effect was more modest than that observed for m^7^GTP (Fig 6B). Surprisingly, m^7^GpppG did not significantly reduce the ability of L375 to decap mRNA, in contrast to the observed reduction of D10 decapping activity by this cap analog (Fig 6C). In support of the specificity of L375 for the methylated cap, the unmethylated versions of the three nucleotides (GDP, GTP, and GpppG) did not significantly inhibit L375 decapping activity. These results suggest that L375 may recognize both the methylated cap structure and the RNA moiety during substrate identification.

**Fig 6.**
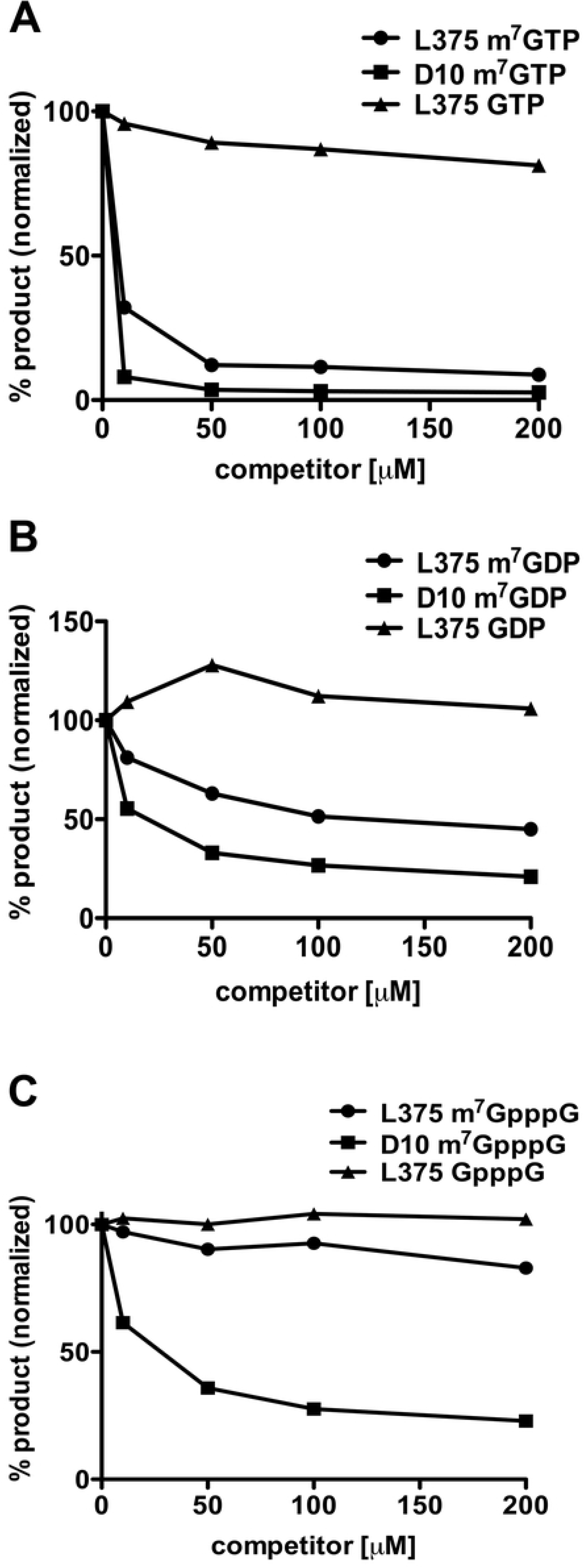
Recombinant L375 mRNA decapping activity is inhibited by addition of m^7^GTP or m^7^GDP. (A) 80 ng of recombinant L375 or VACV D10 and 0.02 pmol ^32^P-cap-labeled actin RNA were incubated together in the presence of increasing quantities of m^7^GTP or GTP in mRNA decapping assays as described in Fig 1B. The percentage of m^7^GDP liberated was calculated through PhosphorImager analysis. (B) mRNA decapping assays were performed as in Panel A with increasing amounts of m^7^GDP or GDP. (C) mRNA decapping assays were performed as described in Panel A with increasing amounts of m^7^GpppG cap analog or GpppG.

## Discussion

Nudix hydrolases cleave a broad range of substrates, a subset of which cleave the 5’ mRNA cap. The importance of these enzymes is illustrated by their conservation between the living organisms (prokaryotes and eukaryotes), as well as their presence in some of the non-living viruses. Three NCLDV Nudix hydrolases have been characterized to date, VACV D9, VACV D10, and ASFV-DP. Each of these enzymes have been shown to possess intrinsic mRNA decapping activity *in vitro* [13–15]. Importantly, ASFV-DP shares sequence similarity to both the well-characterized eukaryotic Dcp2 Nudix mRNA decapping enzyme and Mimivirus L375, the focus of this work [12, 21–26].

Here we show that the Mimivirus Nudix enzyme L375 hydrolyzed the mRNA cap to release m^7^GDP; a reaction that is dependent on an intact Nudix box. Like L375, VACV D9 was shown to preferentially hydrolyze methylated rather than unmethylated mRNA caps [14]. In contrast, ASFV-DP has a broader nucleotide substrate range that includes GTP [15, 20]. Eukaryotic Dcp2 has been reported to preferentially cleave methylated rather than unmethylated cap structures, although some decapping activity has been detected with unmethylated mRNA cap substrates depending on the divalent cation present in the reaction buffer [23–25, 27, 32].

Similar to the viral and eukaryotic mRNA decapping enzymes characterized to date, uncapped competitor RNA reduced L375 decapping activity, suggesting that L375 also uses the RNA moiety for substrate selection [13–15, 25, 27, 33]. Previous studies showed that uncapped competitor RNA more potently inhibited the mRNA decapping activity of ASFV-DP compared to VACV D9 or D10 (ASFV-DP>D9>D10), indicating that ASFV-DP may have a higher affinity for RNA than the poxvirus enzymes [13–15]. In this work, the inhibition of L375 by competitor RNA more closely resembled that of VACV D10, suggesting that L375 may have a lower affinity for RNA than ASFV-DP, despite their greater sequence similarity.

The strong affinity of ASFV-DP for RNA was confirmed through an electrophoretic mobility shift assay, in which the migration of uncapped RNA was impeded with the addition of ASFV-DP [15]. Subsequent *in* vivo experiments confirmed that ASFV interacts with viral and host mRNAs during infection, a property dependent on the N-terminus of the protein, rather than the Nudix motif [21]. ASFV-DP also exhibited different levels of binding efficiency depending on the specific mRNA, similar to Dcp2 [21]. For example, yeast Dcp2 preferentially binds capped mRNAs with a stem-loop structure within the first 10 bases of the sequence. [34, 35].

As observed for VACV D9 and D10 (but not for ASFV-DP), addition of methylated cap derivatives reduced L375 decapping activity, suggesting that L375 recognizes the cap structure in addition to the RNA body [13, 14]. Specifically, L375 decapping activity was most potently inhibited by m^7^GTP, followed by m^7^GDP, whereas the effect of m^7^GpppG was negligible (m^7^GTP>m^7^GDP>m^7^GpppG). In comparison, the decapping activity of both VACV D9 and D10 was diminished by all three methylated cap derivatives, with the most striking reduction also observed with m^7^GTP [13, 14]. Interestingly, VACV D10 was more sensitive to inhibition by methylated nucleotides than either L375 or VACV D9, suggesting a greater importance of the cap structure for substrate recognition for this enzyme [13, 14].

In contrast, ASFV-DP was not inhibited by addition of methylated cap derivatives, nor could it cleave free methylated cap analog *in vitro*, suggesting that for ASFV-DP, the RNA body is critical during substrate identification [15, 20]. Like ASFV-DP, the mRNA decapping activity of most eukaryotic Dcp2 enzymes is not affected by the addition of methylated cap analogs, suggesting these enzymes primarily recognize the RNA moiety to locate target substrates [23–25, 27]. Hence, the inhibition of L375 decapping activity by methylated cap derivatives more closely resembles VACV D9 and D10 rather than that of ASFV-DP or Dcp2.

The purpose of mRNA decapping during viral infection has been revealed through *in vivo* studies characterizing VACV D9, D10, and ASFV-DP. Increased expression of VACV D9, VACV D10, or ASFV-DP resulted in enhanced mRNA turnover of capped viral and host transcripts, further validating that these enzymes decap mRNA to elicit subsequent mRNA degradation [17, 21]. Interestingly, over-expression of ASFV-DP increased degradation of some mRNA transcripts more than others, suggesting that the degree of mRNA turnover induced by ASFV-DP could be selective, as has been shown for Dcp2 [21, 34. 35].

In complementary genetic studies in VACV, deletion or inactivation of D10 resulted in persistence of viral and host mRNAs, a delay in the shutoff of host protein synthesis, and a modest reduction of virulence in animal hosts, whereas deletion of D9 did not produce any noticeable defects [17–19, 36]. When both D9 and D10 were inactivated concurrently, viral replication was severely impaired both *in vitro* and *in vivo*, suggesting that these two enzymes work together synergistically [37]. The substantial replication defects exhibited by the double D9/D10 mutant virus were shown to result from the accumulation of excess amounts of viral double-stranded RNA (dsRNA) that subsequently activated the host’s innate immune defenses to curtail infection [37]. These results suggest that in addition to modulating viral and cellular mRNA turnover, D9 and D10 also induce viral dsRNA degradation to avoid activating host anti-viral responses. Furthermore, cellular RNA exonuclease Xrn1, which degrades RNA following decapping, was also required to avoid aberrant viral dsRNA accumulation, suggesting the viral mRNA decapping enzymes work in concert with cellular Xrn1 to degrade viral dsRNA to evade the host immune system [38].

Like VACV and ASFV, Mimivirus encodes its own mRNA capping enzyme and directs a sequential cascade of viral gene expression consisting of early, intermediate, and late phases; therefore, decapping of viral mRNAs by L375 could provide a mechanism to orchestrate the transitions between viral gene expression during infection [2, 39].

Furthermore, mRNA decapping by L375 during Mimivirus infection could be responsible for the degradation of host mRNAs to elicit the shutdown of host protein synthesis, allowing viral transcripts preferential access to the host translation machinery [2].

In contrast to the animal hosts of VACV and ASFV, Mimivirus primarily infects unicellular *Acanthamoeba* species, and therefore does not require L375 mRNA decapping activity to evade the complex vertebrate immune systems. However, Mimivirus could still use selective mRNA decapping and degradation of specific host mRNAs to promote a robust *Acanthamoeba* infection. Future *in vivo* studies will help elucidate the role of L375 mRNA decapping and decay during Mimivirus infection.

## Acknowledgments

We sincerely thank Bernard Moss for his generous support of this work, guidance, and helpful suggestions. In addition, we are grateful to Wolfgang Resch, P.S. Satheshkumar, Zhilong Yang, and Cheng Huang for helpful discussions and technical assistance.

